# Comprehensive antibiotic-linked mutation assessment by Resistance Mutation Sequencing (RM-seq)

**DOI:** 10.1101/257915

**Authors:** Romain Guérillot, Lucy Li, Sarah Baines, Brian O. Howden, Mark B. Schultz, Torsten Seemann, Ian Monk, Sacha J. Pidot, Wei Gao, Stefano Giulieri, Anders Gonçalves da Silva, Anthony D’Agata, Takehiro Tomita, Anton Y. Peleg, Timothy P. Stinear, Benjamin P. Howden

## Abstract

Acquired mutations are a major mechanism of bacterial antibiotic resistance generation and dissemination, and can arise during treatment of infections. Early detection of sub-populations of resistant bacteria harbouring defined resistance mutations could prevent inappropriate antibiotic prescription. Here we present RM-seq, a new amplicon-based DNA sequencing workflow based on single molecule barcoding coupled with deep-sequencing that enables the high-throughput characterisation and sensitive detection of resistance mutations from complex mixed populations of bacteria. We show that RM-seq reduces both background sequencing noise and PCR amplification bias and allows highly sensitive identification and accurate quantification of antibiotic resistant sub-populations, with relative allele frequencies as low as 10^-4^. We applied RM-seq to identify and quantify rifampicin resistance mutations in *Staphylococcus aureus* using pools of 10,000 *in vitro* selected clones and identified a large number of previously unknown resistance-associated mutations. Targeted mutagenesis and phenotypic resistance testing was used to validate the technique and demonstrate that RM-seq can be used to link subsets of mutations with clinical resistance breakpoints at high-throughput using large pools of *in vitro* selected resistant clones. Differential analysis of the abundance of resistance mutations after a selection bottleneck detected antimicrobial cross-resistance and collateral sensitivity-conferring mutations. Using a mouse infection model and human clinical samples, we also demonstrate that RM-seq can be effectively applied *in vivo* to track complex mixed populations of *S. aureus* and another major human pathogen, *Mycobacterium tuberculosis* during infections. RM-seq is a powerful new tool to both detect and functionally characterise mutational antibiotic resistance.

## INTRODUCTION

Antimicrobial resistance is on the rise and is responsible for millions of deaths every year (World Health Organization 2014). Bacterial populations consistently and rapidly overcome the challenge imposed by the use of a new antibiotic. Their remarkable ability to quickly develop resistance is due to their capacity to exchange genes and to their high mutation supply rate. Multi-drug resistant bacteria are therefore becoming increasingly prevalent and Drug susceptibility testing (DST) is now central to avoid antibiotic misuse and minimise the risk of inducing the emergence of new resistant clones. Over recent years genomics has become a powerful tool to understand, combat and control the rise of resistance (Koser et al. 2014; Schurch and van Schaik 2017). Nevertheless, a precise definition of resistance at the genomic level is crucial to enable fast, culture independent DST by high-throughput sequencing in the clinical context and to track and fight the spread and persistence of resistant clones globally (Van Belkum and Dunne 2013; Schurch and van Schaik 2017).

The genomic basis of resistance is relatively straightforward to establish for resistance conferred by acquisition of a specific gene. The repertoire of resistance genes (resistome) is now well defined and there are several curated databases and software prediction tools for resistance genes detection (McArthur et al. 2013; de Man and Limbago 2016; Liu and Pop 2009). In contrast, comprehensive lists of mutations that confer antibiotic resistance are lacking, despite equivalent clinical relevance. Resistance to major classes of antimicrobials including quinolones, beta-lactams, rifamycins, aminoglycosides, macrolides, sulphonamides, polymyxins, glycopeptides and lipopeptides can all occur via mutations. In some species such as *Mycobacterium tuberculosis*, resistance to all therapeutic agents is mediated by mutations (Smith et al. 2013).

Resistance mutations can be effectively selected *in vitro*, and so genome sequence comparisons of resistant clones derived from sensitive ancestral clones after antibiotic exposure have permitted the identification of numerous resistance-associated mutations (Feng et al. 2009; Livermore et al. 2015; Chen et al. 2014; Mwangi et al. 2007). From these studies it is apparent that the mutational landscape for a single antibiotic combination within a specific bacterium can be broad (Howden et al. 2014; Barbosa et al. 2017; Howden et al. 2010; Handel et al. 2014). Therefore, standard approaches relying on sequence comparisons of single pairs of isogenic mutants are not practical to extensively define the mutational resistome.

Resistance mutations commonly arise in genes encoding the primary drug target or central regulatory genes, such as *gyrA, parC, rpsL, gidB, rpoB, 23S rRNA, rplC, rplD*, and *walKR* (for quinolone, aminoglycoside, rifampicin, linezolid and glycopeptide resistance) (Hershberg 2017; Handel et al. 2014). Because of their implications in central cell processes, such as DNA replication, translation, transcription and cell-wall metabolism regulation, mutations arising in these genes have been associated with a broad range of pleiotropic effects in addition to the antibiotic resistance that they cause (Hershberg 2017). An increasing body of literature shows that antibiotic resistance mutations can lead to broader negative therapeutic consequences through cross-resistance to other antimicrobials (Rodriguez De Evgrafov et al. 2015; Jugheli et al. 2009; Sacco et al. 2015), increased biofilm formation (Yu et al. 2005), increased virulence (Helms et al. 2004; Smani et al. 2012; Beceiro et al. 2013; Gao et al. 2013) and enhanced immune evasion (Gao et al. 2013; Bæk et al. 2015; Cameron et al. 2011; Miskinyte and Gordo 2013). However, there is currently no efficient method to identify pleiotropic mutations. Comprehensively identifying mutations associated with antibiotic cross-resistance and increased risk of therapeutic failure will provide crucial information for future personalised medicine and will help to improve therapeutics guidelines through a greater understanding of the drivers and consequences of mutational resistance. At an epidemiological and evolutionary level, understanding why specific resistance mutations are preferentially selected might provide a rational basis for development of effective measures to combat the rise of resistance.

In this study we developed an innovative workflow called Resistance Mutation sequencing (RM-seq) that enables the unbiased quantification of resistance alleles from complex *in vitro* derived resistant clone libraries, selectable under any experimental condition, allowing identification and characterisation of mutational resistance and its consequences. Here we investigated mutational resistance in *S. aureus* and *M. tuberculosis* and demonstrate that complex resistant sub-populations can be effectively characterised *in vitro* or detected *in vivo* using RM-seq.

## RESULTS

### The RM-seq workflow

RM-seq is an amplicon-based, deep-sequencing technique founded on the single molecule barcoding method (Faith et al. 2013). Here we have adapted this approach in order to identify and quantify at high-throughput, mutations that confer resistance to a given antibiotic (Faith et al. 2013; Kivioja et al. 2011). RM-seq can take advantage of the ability of bacteria to quickly develop resistance *in vitro* to identify and functionally characterise resistance associated mutations at high-throughput. A large and genetically diverse population of resistant clones that encompass the mutational landscape of resistance is selected (Fig 1A). In order to maximise the genetic diversity, a large number of resistant clones (~10,000) are pooled from multiple independent culture and genomic DNA of the mixed resistant population is extracted and the mutational repertoire interrogated by amplicon deep-sequencing.

**Fig 1:**
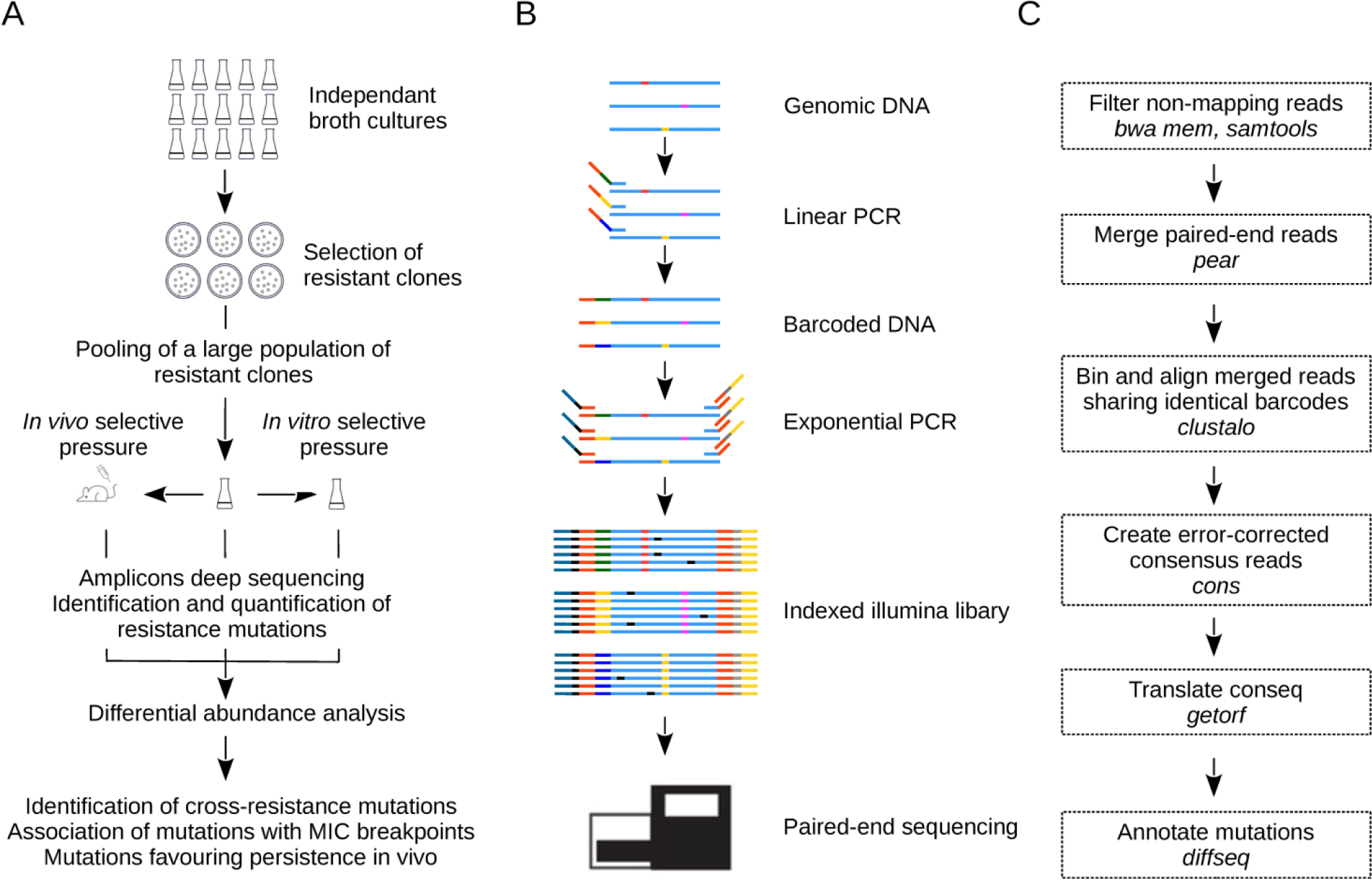
RM-seq workflow. **A.** Schematic view of the experimental design. A large population of resistant clones are selected in vitro from multiple independent cultures. The mutation repertoire selected in a resistance associated locus is then identified by amplicon deep-sequencing. Analysis of the differential abundance of resistance mutations among a resistant clone library before and after a subsequent in vitro (cross-resistance) or in vivo (mouse infection model) selection pressure permits the screening of pleiotropic resistance mutations. **B.** Amplicon library preparation and deep-sequencing. Unique molecular barcodes are introduced by linear PCR (template elongation) using a primer comprising a 16 bp random sequence (green, yellow and blue part of the middle section of the linear PCR primer). Nested exponential PCR using three primers adds Illumina adapters (blue and yellow primer tails) and indices for multiplexing (black and grey primer sections). Grouping of the reads sharing identical 16 bp barcodes allows differentiation of true SNPs (red, pink and yellow) from sequencing errors (black) by consensus sequence reconstruction using multiple reads from the initial template molecule. Counting the number of unique barcodes for each variant provides an unbiased relative quantification of sequence variants. **C.** Bioinformatics analysis pipeline. The diagram represents the different steps in the data processing pipeline. The bioinformatics programs used in the pipeline are indicated in italics.

The high sensitivity and the accurate quantification of the frequency of all the selected mutations in a given genetic loci, enabled screening of complex, mixed libraries of resistant clones. In theory, genetic interactions can be tracked and associated with any selectable pleiotropic phenotype of interest (e.g. cross-resistance to other antimicrobials, immune evasion) by measuring the relative abundance of resistant clones before and after selection. Specific mutations that favour the growth or survival under *in vitro* or *in vivo* test condition will increase in frequency within the population and be readily detected by RM-seq.

Unbiased allele quantification and a low error rate are enabled by single molecule barcoding during the PCR amplicon library preparation (Fig 1B). Sequencing reads sharing identical barcodes are grouped to create consensus sequences of the genetic variants initially present in the population. The single molecule barcoding step has two major advantages. Firstly, it allows error correction of the sequenced DNA and thus high confidence in calling of a resistance associated mutations that might occur at a frequency well below the inherent error rate (~1% (Schirmer et al. 2015)) of the sequencer. Secondly, it permits accurate quantification of allele frequencies by correcting for the amplification bias introduced during the exponential PCR step. The RM-seq bioinformatics pipeline takes as input the raw reads and outputs a table of all annotated substitutions, insertions and deletions identified in the selected population given the original sequence (the target locus sequence before selection). A diagram of the steps in the data analysis pipeline is presented in Fig 1C (RM-seq analysis tool is available from https://github.com/rguerillot/RM-seq).

### Sensitive and quantitative detection of single nucleotide variants in complex bacterial populations

To assess the capability of the RM-seq protocol to detect and quantify rare genetic variants from mixed populations of resistant bacteria, we first evaluated its error correction efficiency. We sequenced at high depth a 270 bp region comprising the rifampicin resistance determining region (RRDR) of a *S. aureus* rifampicin susceptible isolate (wild-type strain NRS384). By counting incorrect nucleotide calls at each position after aligning raw reads to the WT sequence, we found an average error rate per position of 2.8 × 10^−2^ ± 1.7 × 10^−2^ (standard deviation [SD]), which is commonly observed for the Miseq instrument (Schirmer et al. 2015). Merging forward and reverse reads reduced the error rate by an order of magnitude to 5.6 × 10^−3^ ± 4.4 × 10^−3^ (SD). By reconstructing consensus reads supported by at least 10 reads, the RM-seq further reduced the error rate by three orders of magnitude to 1.16 × 10^−5^ ± 3.1 × 10^−5^ (SD) (Fig 2A). At the protein level no further mutations were observed among the 16,516 consensus reads generated (error rate < 6 × 10^−5^).

**Fig 2:**
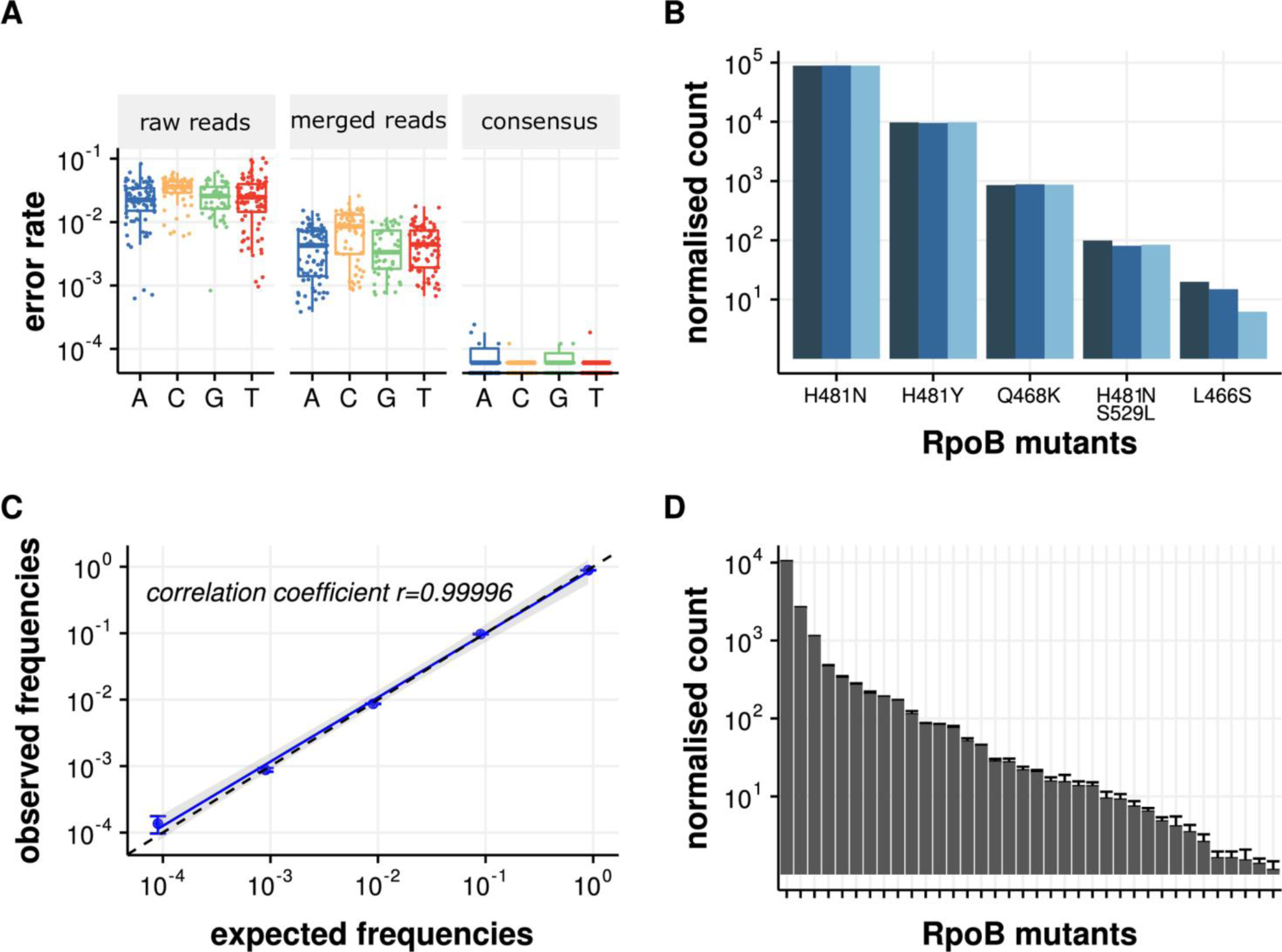
Assessments of the RM-seq protocol. **A.** Error-correction evaluation. RM-seq error-correction combining merging of paired-end reads with consensus sequence determination from grouped reads sharing identical barcode allows a three order of magnitude reduction in false SNP calling when compared with raw reads calling for the different base. **B.** Quantification of populations of S. aureus rpoB mutants. Three independent assessments of rpoB mutants from three independent genomic DNA preparations originating from a defined population are presented by the different blue bars (technical replicates). **C.** Correlation of observed versus expected SNV frequencies. Blue points represent means and error bars represent SEM of three technical replicates. The blue line represents the linear regression of the frequencies measured by RM-seq and the dashed line represent the perfect correlation between expected and observed frequencies. **D.** Quantification of S. aureus rpoB mutants from a complex population of in vitro selected rifampicin resistant mutants. Columns represent mean normalised counts of the different rpoB mutations that were observed among all triplicates, and error bars represent SEM.

We then tested the performance of RM-seq genetic variant quantification on a defined population of genetically reconstructed rifampicin resistant clones. Six different double or single nucleotide variants (SNV) representing different rifampicin resistant *rpoB* mutants were mixed at a relative CFU frequency of 0.9, 0.09, 0.009, up to 0.000009. We applied RM-seq protocol three times independently from three different genomic DNA extractions obtained from this mock community. After library preparation and sequencing on the Illumina MiSeq platform, we obtained 1.8 - 2.2 million raw reads per library, which yielded between 32,433 and 35,496 error-corrected consensus reads, supported by 10 reads or more. At this sequencing depth the mutants ranging from a relative frequency of ~1 to 10^−4^ were readily identified in all three replicates. The normalised count of the different mutants showed little variation between the replicate experiments (Fig 2B) and we observed a very good correlation between the expected mutant frequencies and the observed frequencies after RM-seq (Fig 2C). We also assessed the technical variability of the detection and quantification of RM-seq by independently processing three times the same complex population of *in vitro* selected resistant clones (~10,000 colonies). The relative standard error (RSE) of variant quantification ranged from 0.3% for the most frequent to 38% for rarest variants and the median RSE was 11% (Fig 2D).

### High-throughput identification of rifampicin resistance mutations

In order to comprehensively characterise the mutational repertoire associated with rifampicin resistance we applied RM-seq on the RRDR of three independent pools of ~10,000 colonies capable of growing on agar supplemented with 0.06 mg/L of rifampicin (European Committee on Antimicrobial Susceptibility Testing [EUCAST] non-susceptibility clinical breakpoint). In total, we identified 72 different predicted protein variants; among these 34 were identified in the three independent resistant populations, 17 variants were identified among two resistant populations, and 21 were identified in a single selection experiment (Fig 3). According to our recent extensive literature review of the alleles previously associated with rifampicin resistance (Guerillot et al. 2018), 30 mutations were previously associated with rifampicin nonsusceptibility and 42 alleles identified by RM-seq represent new associations.

**Fig 3:**
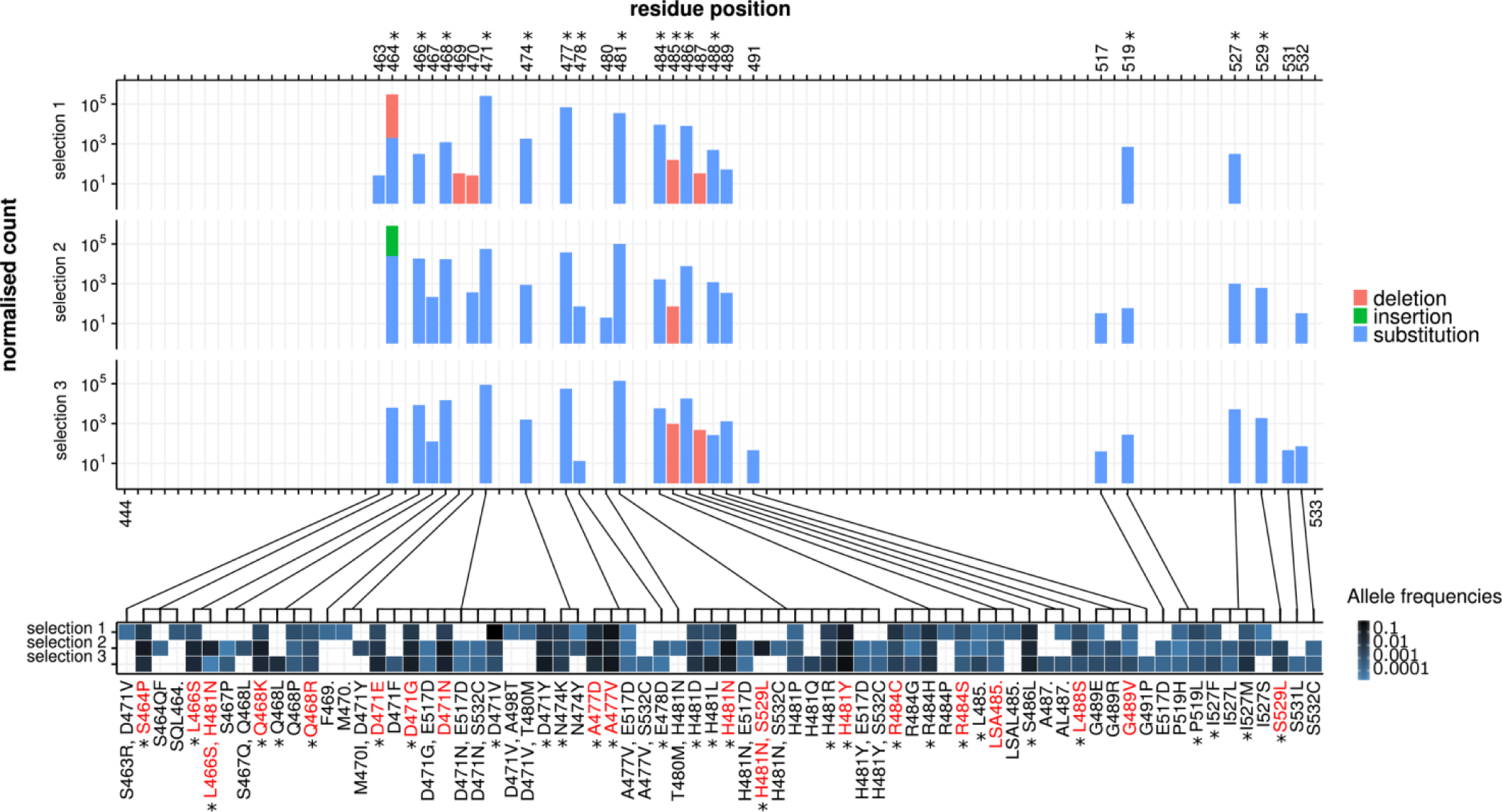
Rifampicin resistance associated mutations detected by RM-seq. Three independent selection experiments of ~10,000 resistant colonies were assessed by RM-seq of the rpoB gene RRDR region. The histograms (upper) represent the normalised mutation counts identified along the sequenced region of the RRDR for the three different selection experiments, with bar colour representing the types of mutation (red for deletions, green for insertions and blue for substitutions). The range of mutations affecting each residue is depicted in the associated heat map (lower panel). The intensity of the blue represents allele frequencies for each selection experiment. Mutations observed from consensus reads reconstructed with at least 10 reads and with a relative frequency greater than 6 × 10^−5^ or identified from all three independent selection experimen ts are represen ted. Resistance mutations that were confirmed by genetic reconstruction are indicated in red (Supplemental_Table_S2.pdf). Mutations and positions previously associated with rifampicin resistance are indicated with a star.

By looking at the different mutated positions, 21 amino-acid positions were repeatedly affected along the RRDR with a similar pattern of mutation frequency at these positions. We observed that 11 different amino acid positions have never previously been associated with rifampicin resistance. The 3D structure modelling of *S. aureus* RpoB protein from *E. coli* RpoB-rifampicin structure showed that the mutated positions were all in close proximity (≤10 Å) to the rifampicin binding pocket of the beta-subunit of the RNA polymerase (Supplemental_Fig_S1.pdf). Therefore, amino-acid sequence alteration at these positions are likely to reduce the rifampicin-RNA polymerase affinity and thus to promote resistance. Interestingly several residues in close proximity to the rifampicin binding pocket were never affected, suggesting that amino acid substitution at these locations do not impair rifampicin binding or that functional constraints make changes to these positions lethal for *S. aureus*. The vast majority of the variants led to amino acid substitutions and several positions, such as 471 and 481 were found to be affected by a high number of different substitutions (11 and 12 respectively). We also observed one complex insertion (S464QF) and eight different deletions.

Positions 485 and 487 represented deletion hot-spots, as they were affected by single, triple and quadruple residue deletions (L485., LSA485., LSAL485.) and single and double deletions (A487. and AL487.), respectively.

We used allelic exchange and site-directed mutagenesis in the WT susceptible background (rifampicin MIC 0.012 mg/L) to reconstruct 19 different *rpoB* alleles that were identified by RM-seq. After whole genome sequencing was used to ensure no secondary non-synonymous mutations or insertion/deletion were introduced (Supplemental_Table_S1.pdf), we confirmed that all these mutations resulted in rifampicin non-susceptibility or resistance with rifampicin MICs above 0.095 mg/L (Supplemental_Table_S2.pdf).

### High-throughput genotype to phenotype associations of resistance mutations with clinical breakpoints

To test if RM-seq could be applied to link a repertoire of resistance mutations to a particular resistance threshold, we selected rifampicin resistant clones, grown on plates supplemented with different concentrations of antibiotic (in this case, rifampicin). To select resistant subpopulations we used the most widely used clinical resistance breakpoints from the guidelines of the EUCAST and the Clinical & Laboratory Standards Institute (CLSI) (EUCAST 2015; CLSI. Performance standards for antimicrobial susceptibility testing 2016). Therefore, we selected sub-populations growing on plates supplemented with rifampicin at concentrations of 0.06 mg/L (EUCAST non-susceptibility), 0.5 mg/L (EUCAST resistance), 1 mg/L (CLSI nonsusceptibility) and 4 mg/L (CLSI resistance). The result of resistant sub-population detection and quantification by RM-seq associated with the different antibiotic concentration thresholds is presented in Fig 4. Among 43 mutations, 24 mutations were detected at all antibiotic concentration thresholds and therefore would be classified as resistance-conferring mutations by both guidelines. Among the 19 other mutations detected, four were associated with resistance levels ranging from 1 to 4 mg/L, three with resistance ranging from 0.5 to 1 mg/L and the remaining 12 with resistance ranging from 0.006 to 0.5 mg/L. Interestingly, *S. aureus* with any of these last 12 alleles, selected only at low antibiotic concentrations, would be classified as non-susceptible by EUCAST and susceptible by CLSI. Similarly, *S. aureus* with three mutations associated with resistance by EUCAST would be classified as susceptible by CLSI (Fig 4).

**Fig 4:**
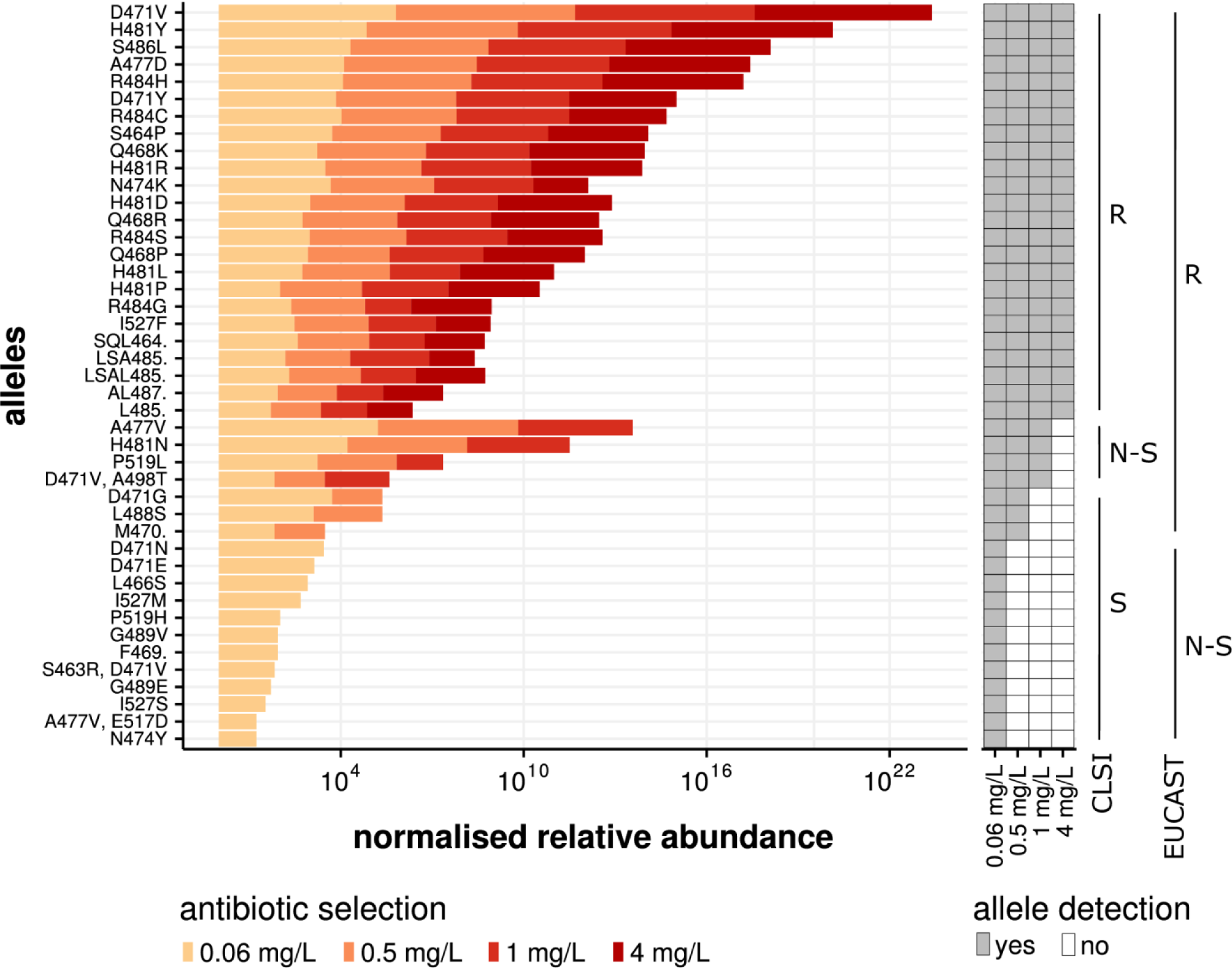
Association of resistance mutations with clinical MIC breakpoints. The histogram represents the relative abundance of individual mutations recovered from the selected subpopulation. The colour yellow to red represents the rifampicin concentration used for selection. The antibiotic concentrations were chosen according to the CLSI and EUCAST guidelines (see legend). The detection (grey box) and disappearance (white box) of a particular allele from the population at the different antibiotic selection break-points is depicted on the right of the histogram. The presence or absence of allele detection at the different antibiotic concentration breakpoints were used to associate the alleles with sensitive, non-susceptible or resistant classification of the CLSI and EUCAST guidelines (S, susceptible; R, resistant; N-S, non-susceptible).

We used mutants reconstructed by allelic-exchange to verify that the resistance level predicted by RM-seq matched the MIC conferred by a particular allele. Among 17 reconstructed mutants tested, 16 showed MICs in complete accord with the RM-seq prediction (Supplemental_Table_S2.pdf). One mutant (D471G) with a borderline measured MIC of 0.5 mg/L was predicted to have an MIC superior to 0.5 and inferior or equal to 1 despite showing clear reduction in abundance on 0.5 mg/L plate by RM-seq (Fig 4).

### High-throughput screening of resistance mutations associated with antimicrobial crossresistance or collateral sensitivity

In order to evaluate if RM-seq can be used to characterise pleiotropic resistance mutations that confer an increased or decreased susceptibility to a second antibiotic (cross-resistance or collateral sensitivity respectively), we followed the differential abundance of resistance mutations of a complex rifampicin resistant population after selection with a second antibiotic, daptomycin. We chose daptomycin because it is a last-line antibiotic used against multidrug resistant *S. aureus*, commonly deployed in combination therapy with rifampicin to treat complicated infections (Forrest and Tamura 2010; Saleh-Mghir et al. 2011; Garrigos et al. 2010). Furthermore, some *rpoB* mutations have been previously associated with subtle changes in daptomycin MIC (Cui et al. 2010; Aiba et al. 2013). We screened for pleiotropic effects on daptomycin resistance by performing three independent time killing experiments using a large *in vitro* derived population of rifampicin resistant clones. Daptomycin concentrations of 8 mg/L corresponding to the minimal plasma concentration commonly reached during standard antibiotic therapy were used (Reiber et al. 2015). Survival of the rifampicin resistant population at 3 hours represented 1.6% (± 0.1 SEM) of the initial inoculum and bacterial regrowth was observed to 8.8% (± 6.9 SEM) at 24 hours (Fig 5A). The abundance of all rifampicin resistance mutations were then quantified by RM-seq for the initial bacterial population (the inoculum) and the surviving population at three hours and 24 hours after daptomycin exposure for the three independent killing experiments (Fig 5B).

**Fig 5:**
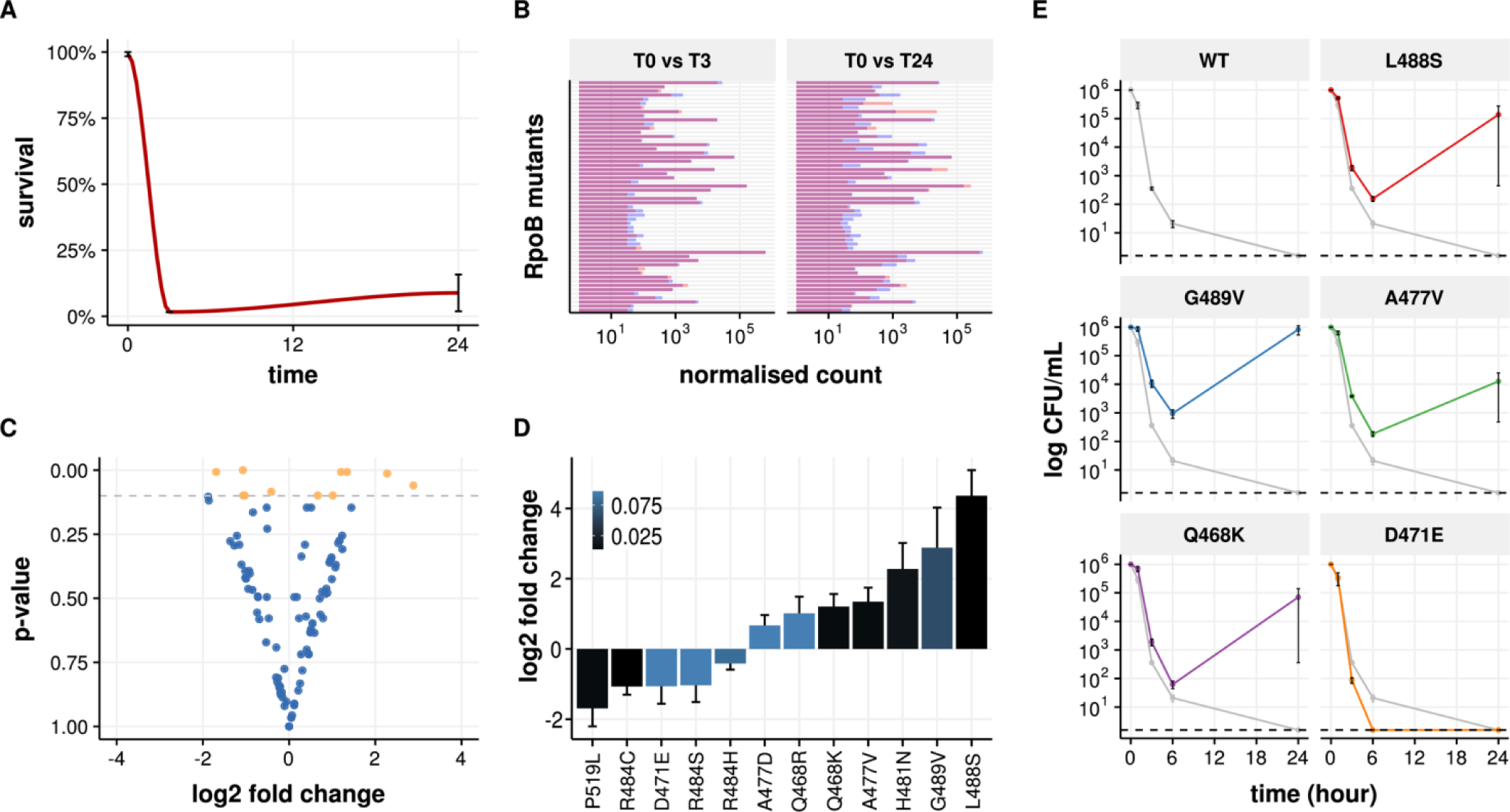
Screening of resistance mutations associated with cross-resistance or collateral sensitivity. **A.** Daptomycin selection (8 mg/L) of a pooled population of in vitro selected rifampicin resistant clones. Survival was quantified by CFU counting on BHI agar plates at 3 hours and 24 hours of exposure. Error bars represent ± SEM of three independent exposures to daptomycin. **B.** Rifampicin resistant mutant quantification of rifampicin mutant before and after 3 hours or 24 hours of daptomycin exposure. Each bar of the histogram represents the averaged normalised count of the different rpoB mutants in the population. Average quantification of the three replicates at T=0 and after daptomycin exposure are indicated by blue and red bars respectively. Bars are superimposed for each mutant and overlap of the bars are coloured in purple. Increases and decreases in allele frequencies after daptomycin exposure are indicated by red and blue bars respectively on the top of purple bars. **C.** Volcano plot showing fold change in rpoB alleles frequency after 24 hours of daptomycin exposure. Each dot represents a different rpoB mutant. Orange dots represent mutants with p-value <0.1 by Wald test. **D.** Rifampicin resistance mutations associated with significant fold change after 24 hours of daptomycin treatment. Mutations with positive and negative log2 fold change are predicted to be associated with cross-resistance and collateral sensitivity to daptomycin, respectively. The intensity of the blue coloration of the bars represents adjusted p-values (Wald test). **E.** Daptomycin time kill assays. Rifampicin resistant mutants were assessed in triplicates (biological replicates), points represent the mean survival at each time point and error bars SD. Dashed lines represent detection limit.

We tested for significant differential abundance of all the different mutations detected (Fig 5C). After 3 hours of daptomycin treatment, one mutation appeared to increase in frequency (Q468K) and another decreased (P519L) but the null hypothesis (no change) could not be rejected (p>0.05 after correction for multiple testing [Wald test]). At 24 hours of daptomycin selection, differential abundance of these two mutations increased, together with 10 other rifampicin resistance mutations when compared with the mutant abundance in the initial population (Fig 5D). All the rifampicin resistance mutations that were previously identified as conferring decreased susceptibility to daptomycin (n=6) were found enriched after daptomycin selection, and four mutations had significant fold changes at the 24-hour time point (p<0.05, Wald test). These experiments show that changes in relative allele abundance as measured by RM-seq are concordant with changes in daptomycin susceptibility (Berti et al. 2015; Aiba et al. 2013).

In order to validate the use of RM-seq as a screening method to identify new mutations that confer cross-resistance or collateral sensitivity we introduced in the wild-type strain by allelic exchange seven rifampicin resistant mutations that were significantly enriched and three mutations that were significantly rarefied after daptomycin selection. Among these mutations, MIC testing validated six of the seven rifampicin resistance mutations as decreasing susceptibility to daptomycin and one mutation as increasing the daptomycin susceptibility (Supplemental_Table_S2.pdf). We then performed daptomycin time kill assays and found that even though the D471E mutation did not show a decreased MIC to daptomycin (Supplemental_Table_S2.pdf), this mutant was less tolerant to daptomycin (Fig 5E), concordant with the RM-seq prediction which demonstrated reduced abundance of this mutation after daptomycin exposure (Fig 5D). Similarly, rifampicin resistance mutations L488S, G489V, A477V and Q468K were clearly associated with increased tolerance to daptomycin killing (Fig 5E).

Taken together our data demonstrate that RM-seq can identify pleiotropic resistance mutations conferring changes in susceptibility to a secondary antibiotic from large pool of resistant clone selected *in vitro* after exposure to a primary antibiotic.

### Tracking resistant clones *in vivo* in a mouse infection model

The relationship between resistance selection, *in vivo* fitness cost and pathogenicity has been a long standing research topic (Beceiro et al. 2013; Cameron et al. 2011; Holmes et al. 2011; Gao et al. 2010; Beceiro et al. 2012). In a proof-of-principle experiment to investigate the dynamics and fitness of resistance mutations *in vivo*, we followed the abundance of rifampicin resistance mutation by RM-seq in a mouse model of persistent infection. Six-week-old BALB/c mice were injected via the tail vein with a complex, *in vitro* derived population of rifampicin resistant mutants that also included susceptible WT clones. We then quantified the abundance of RpoB mutants in the inoculum and at 1 and 7 days post-infection in the kidney, liver and spleen of the mice (Fig 6). At 24 hours post infection, we recovered a diverse set of RpoB mutants with different relative abundances in the two mice tested. The diversity of mutants appeared to be reduced when compared with the inoculum in the different organs and several initially abundant mutants were not recovered showing a rapid clearance of several inoculated clones. Interestingly, at 7 days post-infection we observed a drastic reduction in resistant clone diversity with only a small number of clones dominating. This result supports the concept that the establishment of *S. aureus* infection in the mouse is highly clonal, following a “bottleneck” in which very few bacterial cells establish infectious foci or abscesses in invaded organs (McVicker et al. 2014). Despite the intravenous inoculum containing a diversity of resistant clone, we observed that within a given mouse different organs were infected with the same clones.

**Fig 6:**
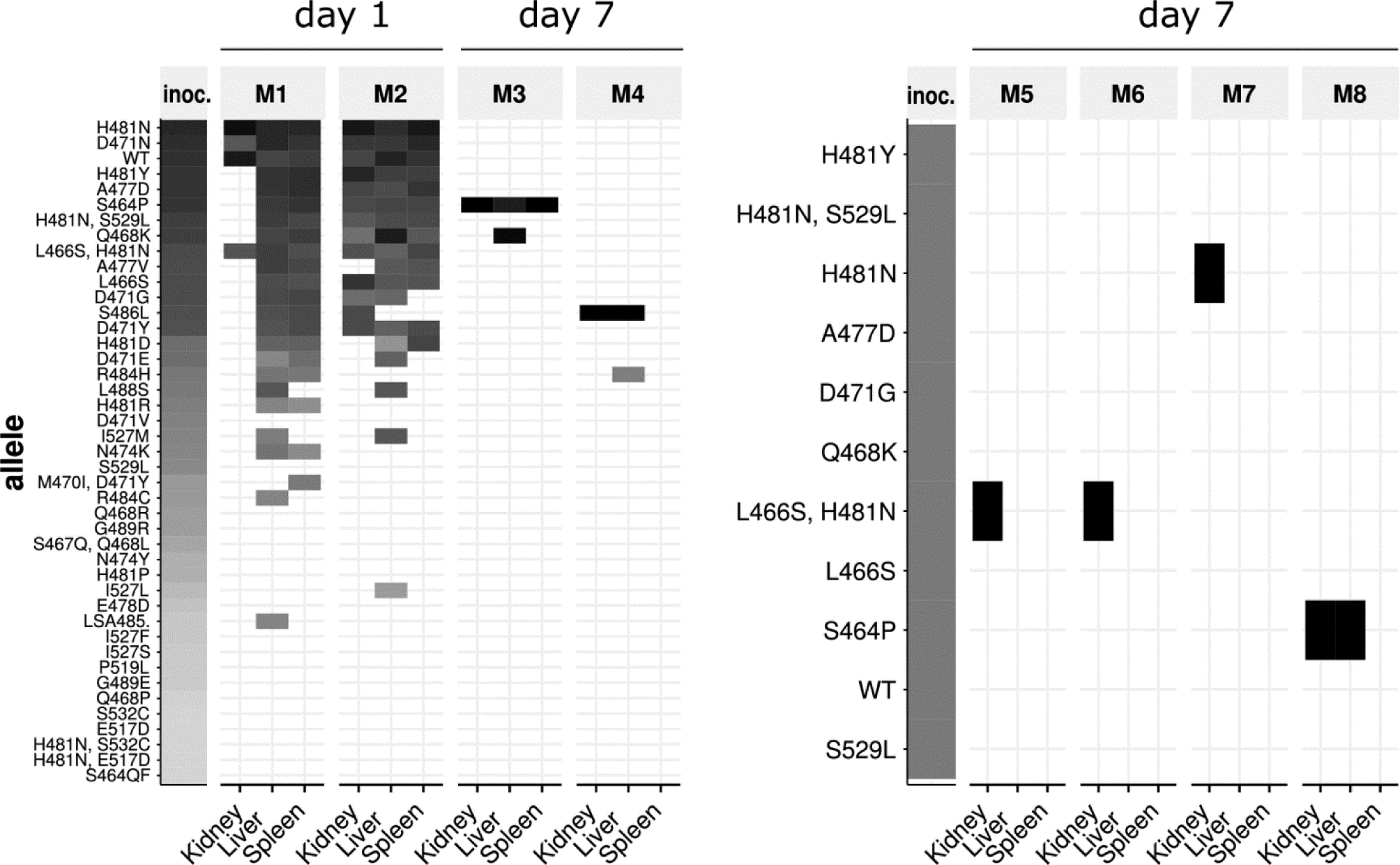
In vivo detection of rifampicin resistance mutations in a mouse persistence model. The heat maps represent quantification of RpoB mutants in kidney, liver and spleen of eight different mice after 1 or 7 days infection with a complex in vitro selected population (mice Ml to M4 on the left) or with a genetically defined population of rifampicin resistant clones (mice M5 to M8). The columns labelled ‘inoc.’ represent the initial inoculum. Grey and black boxes represent low and high relative allele abundance.

We then infected mice with an inoculum comprising an equal amount of 10 reconstructed RpoB mutants together with the wild-type susceptible strain. After 7 days of infection four mice were analysed for RpoB mutant abundance by RM-seq. As observed in the previous experiment, the wild-type allele did not persist after 7 days, showing that resistant clones are not outcompeted by the wild-type clone for persistence in the mouse model; even without antibiotic selective pressure (Fig 6). Three out of four mice were infected with clones encoding the H481N mutation, which has been found to be the most frequent mutation among sequenced *S. aureus* human isolates (Guerillot et al. 2018), two had the L466S, H481N double mutation and one had H481N only. Intriguingly, because the mice were infected simultaneously with 11 different clones, the probability is low that at least two mice would become infected with the L466S, H481N by chance (p=0.043). The probability is also low that at least three mice would become randomly infected by a clone encoding the H481N mutation (p=0.057). Given the relatively small number of mice investigated here, no conclusions can be drawn on the potential competitive advantage of specific resistance mutations *in vivo.* Nevertheless, we show here that RM-seq can be used to follow the dynamics of complex populations of clones in a mouse infection model and that the design of complex multi-clone competition assays *in vivo* is achievable with RM-seq.

### Detection of low frequency resistant sub-populations of *M. tuberculosis* from sputum samples

A primary motivation for developing RM-seq is to reduce inappropriate antimicrobial therapy by allowing the early detection of low frequency drug resistant sub-populations that can arise during antimicrobial therapy. Treatment of tuberculosis, caused by infection with *M. tuberculosis*, could be significantly improved by an accurate and sensitive amplicon sequencing method. This is because culture-based methods to detect resistance can take weeks to obtain a result and current rapid molecular diagnostic methods only detect a handful of commonly occurring mutations and have low sensitivity for the detection of resistant subpopulations (Zetola et al. 2014). A technique that was comprehensive and relatively rapid, particularly when infection with multi-drug resistant *M. tuberculosis* was suspected, would arm clinicians with rich data to inform effective antibiotic treatment regimens.

To assess the potential applicability of RM-seq for clinical detection of resistant sub-population we retrospectively applied RM-seq on genomic DNA extracted from sputum samples of a patient affected by chronic pulmonary multi-drug-resistant tuberculosis. Multiple sputum isolates from this case of chronic *M. tuberculosis* infection have been previously investigated by whole genome sequencing (Meumann et al. 2015). Here we investigated the emergence of resistance mutations from two samples (sampling interval of 11 years) of three different loci in the genes *rpoB, pncA* and *ethA* associated with resistance to rifampicin, pyrazinamide and ethionamide. The multiple changes that were made to the treatment regimen are summarized in Fig 7. Using RM-seq we found four other low frequencies *rpoB* mutants in addition to the dominant *rpoB-* S450L alleles previously associated with rifampicin resistance in this case (Fig 7)(Donnabella et al. 1994). Among those, rpoB-H445N (frequency of 1.24 × 10^−3^) and rpoB-G442W (frequency of 6.86 × 10^−4^) represent known rifampicin resistance conferring alleles (Ramaswamy et al. 2004; Pozzi et al. 1999). In the later sputum samples collected 12 years after the end of rifampicin treatment these low frequencies sub-populations of rifampicin resistant clones were not detected but the dominant rifampicin resistant population harbouring mutation *rpoB-* S450L persisted together with a low frequency population harbouring the rpoB-P471Q allele. In association with pyrazinamide resistance the resistant allele pncA-I31F dominated the population after a year and half of treatment together with a low frequency of double mutant sub-population represented by the allele pncA-R29S-I31F. The resistant mutant pncA-I31F was also detected on the later isolate. Surprisingly, early samples were susceptible to pyrazinamide as established by phenotypic testing despite a high prevalence of the pncA-I31F resistant allele. For ethionamide resistance, the wild-type version of the gene *ethA* was initially dominant in the population (frequency of 8 × 10^-1^) together with a low frequency allele not associated with resistance ethA-H102P (frequency of 2 × 10^-1^). The ethionamide resistance mutation ethA-*ΔA338* causing a frameshift in the gene was readily detected in accordance with phenotypic testing in the later sputum sample. Thus, RM-seq was able to identify low frequency subpopulations of antibiotic resistant *M. tuberculosis*.

**Fig 7:**
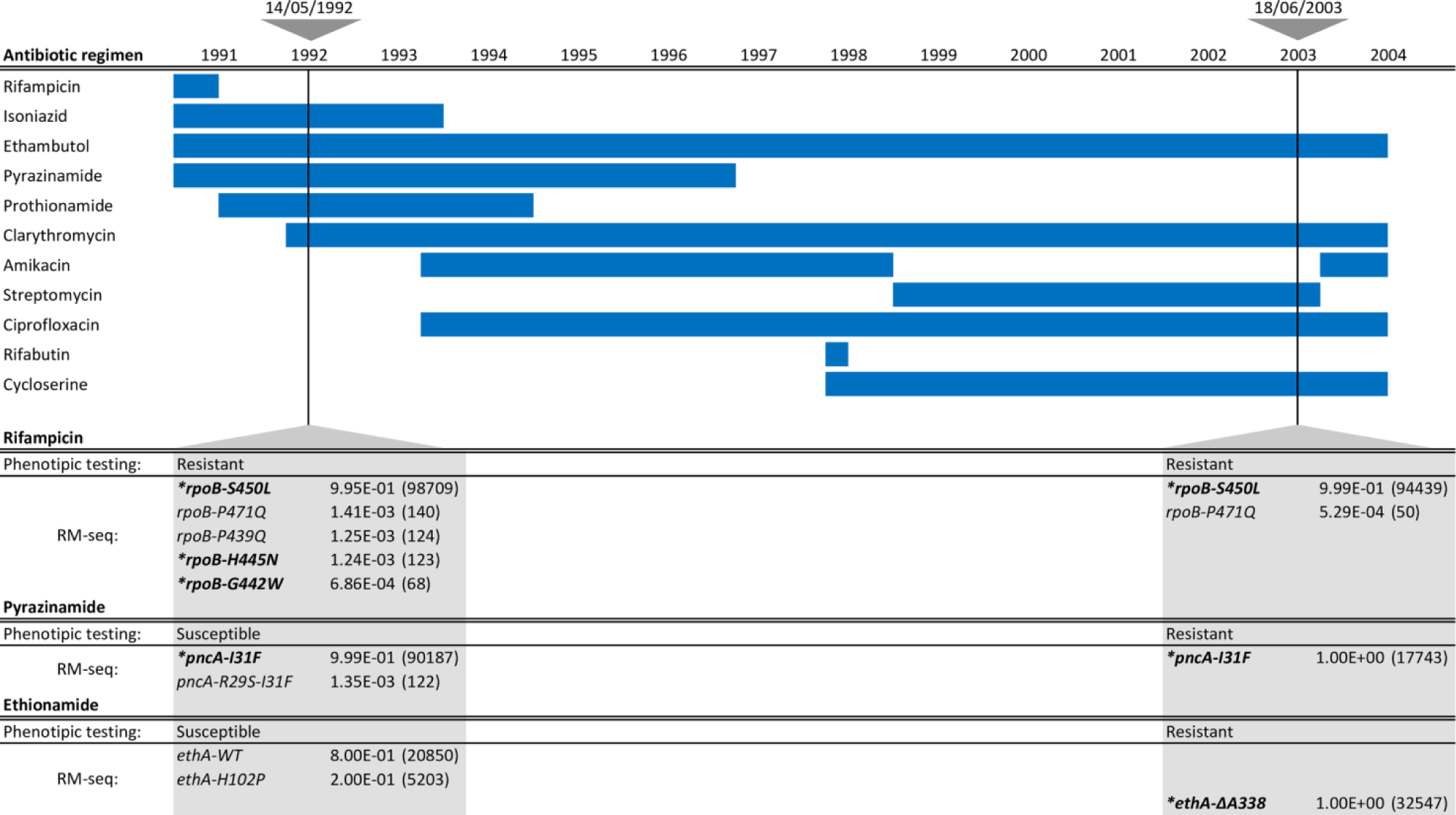
Detection of low frequency resistant sub-populations of M. tuberculosis from sputum samples. Depicted on the top left are the antibiotics used with the date that each treatment was initiated and the duration indicated by the blue horizontal bars. The two triangles at the top of the figure represent the early and late DNA extracts used for RM-seq. The table at the bottom shows phenotypic testing and RM-seq results for rifampicin, pyrazinamide and ethionamide for the two samples tested. For RM-seq results the frequency of each allele is indicated and the number of consensus reads is in parenthesis. Alleles in bold and annotated with star represent alleles known to confer antibiotic resistance. Consensus reads reported were reconstructed from at least six reads and alleles represented by at least 50 consensus reads.

## DISCUSSION

In this study, we designed and validated a new high-throughput workflow call RM-seq that enables fast and comprehensive characterisation of antibiotic resistance mutations. We show that a straightforward molecular PCR-based barcoding step coupled with high throughput sequencing significantly reduces background sequencing noise and permits accurate identification and quantification of rare resistance mutations in complex bacterial populations. By applying RM-seq on large pools of *in vitro* selected rifampicin resistant *S. aureus* clones, we demonstrate that the mutational resistome of a resistance locus can be defined. We found that the range of rifampicin resistance mutations in *S. aureus* is broader than previously understood, highlighting the inadequacy of our understanding of the genetic basis of resistance. Here, 72 mutations were associated with rifampicin resistance in *S. aureus.* In comparison, the CARD database of antibiotic resistance markers only contains 6 rifampicin resistance mutations (McArthur et al. 2013). As the RM-seq protocol can be applied on any combination of microorganisms and resistance, its use has the potential to greatly enhance current knowledge on microbial adaptation to antibiotic exposure.

One limitation of RM-seq is the size limit of the sequenced region that can be interrogated by a single amplicon (270 bp with fully overlapping reads). This limitation is imposed by the maximum read length of Illumina^®^ paired-end sequencing technology. Nevertheless, because RM-seq is compatible with standard Nextera^®^ indexing primers, up to 384 resistance targets can be multiplexed in a single sequencing run. Furthermore, when performing read subsampling simulation, we found that the number of high quality consensus reads increase almost linearly with the number of reads when performing low depth sequencing (Supplemental_Fig_S2.pdf). As little as 140,000 reads would be sufficient to obtain 10,000 consensus reads supported by 10 reads. Theoretically, all resistance variants arising among more than 350 different targeted regions of 270 bp would be accurately identified from a mixed population 1000 bacterial clones using a single MiSeq run (94500bp, ~86 different genes). This kind of experimental design would be valuable to characterise the genetic basis of poorly defined resistance mechanisms or to determine all the resistance mutation arising in a particular gene. During the preparation of this manuscript, we scanned the full genes *mprF* (2523 bp) and *cls2* (1482 bp) for mutations conferring daptomycin resistance in *S. aureus* by sequencing 10 and six amplicons respectively (manuscript in preparation). For the development of diagnostic tool multiple resistance hotspots could be assessed by RM-seq using a similar design.

The application of RM-seq is not restricted to the high-throughput identification of resistance mutations and can also be used to characterise the phenotypic impact of specific resistance mutations. We demonstrate here that differential mutation abundance analysis can be performed to link subsets of mutations with clinical resistance breakpoints and to identify resistance mutations that favour survival or multiplication in particular conditions. Comparisons of allele frequencies in mixed populations before and after exposure to a second antibiotic permitted the identification of specific resistance mutations that confer crossresistance and collateral sensitivity. The demonstration that several specific rifampicin resistance mutations can prevent bacterial clearance by daptomycin *in vitro* can have potential clinical implication regarding the usage rifampicin and daptomycin in combination therapy. A deeper understanding of how evolution of microbial resistance towards a given antibiotic influences susceptibility or resistance to other drugs would have profound impact as it could be exploited to fight resistance rise through combination therapy or by the temporal cycling of different antibiotics (Pal et al. 2015; Rodriguez De Evgrafov et al. 2015; Imamovic and Sommer 2013).

We also showed that the persistence of resistance alleles can be followed during experimental infection (murine blood stream infection model). As it is known that specific resistance mutations can favour pathogenesis and immune evasion (Beceiro et al. 2013; Gao et al. 2013; Bæk et al. 2015), RM-seq can be used to screen for resistance mutations that increase or decrease survival against *ex vivo* selective pressures (eg. whole blood killing, phagocytosis, antimicrobial peptide killing, complement killing) or that favour colonisation or tissue invasion (eg. biofilm formation, cell attachment, intracellular persistence). A better characterisation of critical resistance mutations that confer cross-resistance or that impact pathogenesis would permit both improving antibiotic resistance surveillance and drug management if a higher therapeutic risk is confirmed.

Fast and culture independent molecular diagnostic tools have revolutionised pathogen identification and resistance typing in clinical settings. We show here that RM-seq can be used to detect very low frequency sub-population of resistant clones from patients infected by *M. tuberculosis.* The development of diagnostic tools based on the combination of PCR-based barcoding and massively parallel sequencing represents a promising approach for the next generation of genetic-based diagnostics. RM-seq has potential advantages over standard quantitative and molecular probe-based diagnostic tests. For instance, RM-seq would be more sensitive than the current best practice platform for rifampicin resistance detection in *M. tuberculosis*, GeneXpert, as this platform fails to identify sub-populations of rifampicin-resistant strains representing less than 10% of the population (Zetola et al. 2014) and digital PCR and qPCR assays that have been validated for rare mutations with frequencies-of-occurrence not lower than 0.1% (Whale et al. 2016). This property of RM-seq may have important clinical implications as similarly to molecular test, most phenotypic tests fail to detect heterogeneous resistance with resistance allele frequency below 1%, and lower frequency of resistance have been frequently described (Eilertson et al. 2014; Koser et al. 2014; Howden et al. 2014). RM-seq detection is not conditional on the affinity of short DNA probes, therefore all sensitive and resistant variant can be detected and differentiated at the sequence level. Therefore, diagnostic tools based on molecular barcoding and deep sequencing have the potential to perform better than current state of the art diagnostic tests by accurately detecting pre-existing rare resistant sub-population as well as uncommon resistance mutations.

We expect that RM-seq will be a valuable tool for the comprehensive characterisation of the mutational resistance repertoire. A deeper understanding of resistance at the DNA level will be the basis for improved genomic surveillance of antibiotic resistant pathogens, optimised antibiotic treatment regimens, and can ultimately lead to precision medicine approaches for treating microbial infections.

## METHODS

### *In vitro* selection of rifampicin resistant clones

All experiments were conducted with *S. aureus* USA300 strain NRS384, acquired from BEI resources. Rifampicin resistant colonies were selected from 20 independent overnight Heart Infusion (HI) 10 mL broth cultures (5x10^9^ CFU/mL) inoculated from single colonies. Cultures were pelleted at 10 min at 3000 g and re-suspended in 200 μL of HI broth (2.5x10^n^ CFU/mL). These concentrated overnight cultures were then pooled and plated on HI plates supplemented with rifampicin at 0.006, 0.5, 1 and 4 mg/L. Given that the spontaneous resistance rate for rifampicin in *S. aureus* is ~2x10^−8^ (O’Neill et al. 2001), 20 to 30 plates inoculated with 75 μL were necessary to recover ~10,000 resistant clones after 48h incubation at 37°C. All resistant colonies were recovered by scraping the plate flooded with 2 mL of Phosphate Buffered Saline (PBS). After washing the pooled clone libraries in PBS, aliquots were used for genomic DNA extraction and RM-seq library preparation and stocked in 25% glycerol at -80°C.

### Amplicon library preparation and deep-sequencing

Genomic DNA was extracted from 1 mL aliquots adjusted to an OD600 of 5 in HI broth. Cells were pelleted and washed twice in PBS and genomic DNA was extracted using the DNeasy Blood & Tissue Kit (QIAGEN). Random 16 bp barcodes were introduced by performing 8 cycles of linear PCR with the primer *x_RMseq_F* (Supplemental_Table_S3.pdf) using the following PCR mix: 2 μL of *x_RMseq_F* (5nM), 1 μL of genomic DNA (6 ngμL), 12.5 μL Phusion^®^ High-Fidelity PCR Master Mix (2X, New England BioLabs Inc.), 6 μL H20. The following PCR cycle conditions were used: 30 sec at 98°C, then 8 cycles of 10 sec at 98°C, 30 sec at 50°C, 30 sec 72°C, and a 2 min elongation step at 72°C. Following the final cycle of the linear PCR, samples were cooled to 25°C and the nested exponential PCR were performed by immediately adding 3.5 μL of a primer mix containing 2 μL of primerx *_RMseq_R* (100 nM), 0.6 μL forward and 0.6 μL reverse Nextera XT Index Kit primers (10 μM), 0.3 μL H2O. The PCR conditions above were then used for a further 25 cycles. The resulting amplicons comprising Illumina adaptor and indices was purified with Agencourt^®^ AMPure^®^ XP magnetic beads (Beckman Coulter) using beads/sample volume ratio of 0.8. Purified amplicons were then normalised at 4 nM according to expected size and measured DNA concentrations (Qubit™ dsDNA HS Assay Kit). Amplicons with different indices were pooled and the sequencing library was diluted to 15 pM with 10% *phiX* control spike and sequenced on Illumina Miseq or Nextseq using Reagent Kit v3 to produce 300 bp or 150 bp paired-end reads. Sequencing reads of RM-seq experiments are available from NCBI/ENA/DDBJ under BioProject number PRJNA399605.

### Bioinformatics analysis pipeline

The RM-seq pipeline processes raw reads after demultiplexing by the Illumina sequencing instrument. The pipeline uses *bwa mem* read aligner (0.7.15-r1140) (Li 2013) to map reads to a reference locus *(rpoB)* and *samtools* (v1.3) (Li et al. 2009) to remove unmapped and low quality reads from the read sets. Then *pear* (v0.9.10) (Zhang et al. 2014) is used to merge paired reads. Merged reads sharing identical barcodes are aligned using *Clustal Omega* (v1.2.1) (Sievers and Higgins 2014). Cons from the EMBOSS suite (v6.6.0.0) (Rice et al. 2000) is used to collapse the alignments into single error-corrected consensus reads. To speed-up processing, read alignment and consensus sequence generation tasks are executed in parallel using *GNU parallel* (Tange 2011). Unique consensus DNA sequences are identified via clustering using the *cd-hit-est* module of the *CD-HIT* (v4.7) software (Fu et al. 2012). Resultant unique representative consensus sequences are translated to amino acids using *getorf* and annotated at the protein and nucleotide level using *diffseq*, both modules of the EMBOSS suite. The annotated effect of mutation is then re-associated to each barcode in the final output table.

### Construction of *rpoB* mutants by allelic exchange

Allelic exchange experiments were performed using shuttle vector pIMAY-Z (Monk et al. 2015) with some modifications. Full-length *rpoB* sequences corresponding to the 19 different *rpoB* alleles reconstructed by allelic exchange in the *S. aureus* NRS384 strain were obtained by performing PCR overlap extension with Phusion High-Fidelity DNA Polymerase (New England Biolabs) and introducing *rpoB* codon mutations to the primer tails (Supplemental_Table_S3.pdf). Gel purified *rpoB* amplicons were then joined with pIMAY-Z using Seamless Ligation Cloning Extract (SLiCE) cloning (Zhang et al. 2012) and transformed into *E. coli* strain IM08B (Monk et al. 2015) to allow CC8-like methylation of the plasmid and bypass the *S. aureus* restriction barrier. The presence of a cloned *rpoB* insert in pIMAY-Z plasmid was then confirmed by colony PCR using primers pIMAY-Z-MCSF and pIMAY-Z-MCSR. Purified plasmid was then electroporated into *S. aureus* and plated on HI supplemented with chloramphenicol at 10 mg/L and X-gal (5-bromo-4-chloro-3-indolyl-β-d-galactopyranoside; Melford) at 100 mg/L and grown 48h at 30°C. Blue colonies were picked and grown in HI broth at 37°C without Cm selection pressure overnight to allow loss of the pIMAY-Z thermosensitive plasmid. Double cross-overs leading to allelic replacement of the wild type with the desired rifampicin resistant *rpoB* alleles were directly selected by plating cultures on HI plates supplemented with 0.06 mg/L of rifampicin. Rifampicin resistant and chloramphenicol sensitive colonies arising at a frequency higher than 10^−3^ were considered as potentially positive clones for allelic exchange as spontaneous rifampicin resistance arises at a much lower frequency of ~2x10^-8^ (O’Neill et al. 2001) in the wild type strain. Clones were then colony purified on HI plates before glycerol storage and extraction of genomic DNA. To validate the allelic exchange procedure, the whole genome sequence of all reconstructed strains was determined with the Illumina Miseq or Nextseq 500 platforms, using Nextera XT paired-end libraries (2x300 bp or 2x150 bp respectively). To ensure that no additional mutations were introduced during the allelic exchange procedure, reads of all mutant strains were mapped to the reference NRS384 genome (Monk et al. 2015) using Snippy (v 2.9) (https://github.com/tseemann/snippy). The results of the SNP/indel calling of the reconstructed mutants were then compared with our NRS384 WT reference isolate. The SNP/indel profile for each mutant is presented in Supplemental_Table_S1.pdf.

### Antibiotic susceptibility testing and time kill assays

Rifampicin and daptomycin MIC were measured using E-tests (BioMerieux) on Mueller-Hinton plates supplemented with 50 mg/L Ca^2^+ following manufacturer’s instructions. For daptomycin time kill assays, 10 mL of BHI broth supplemented with 8mg/L daptomycin and 50 mg/L Ca^2^+ was inoculated with 10^6^ CFU/mL of an overnight culture. Cultures were incubated at 37°C with constant shaking and samples were collected at 3 hr, 6 hr and 24 hr time points. Cell survival after daptomycin exposure was assessed by calculating the ratio of the CFU at 3, 6 and 24 hours on the CFU of the initial inoculum (10^6^ CFU/mL) and taking the average colony counts of duplicate BHI agar plates. All daptomycin time kill assays were performed in biological triplicate.

### Mutant differential abundance analysis after daptomycin selection

Three replicates of daptomycin selection were performed on a rifampicin selected population *(in vitro* 1 population selected with rifampicin at 0.06 mg/L). A high initial inoculum of 5x10^8^ CFU/mL was used to recover a sufficient amount of bacterial DNA from surviving cells after daptomycin exposure. After 3 hours or 24 hours of exposure to daptomycin at 8 mg/L, surviving bacterial populations were pelleted and washed. To remove extracellular DNA resulting from daptomycin induced cell death, cell pellets were incubated 45 min at 37°C with 1 μL of Amplification Grade DNase I (1U/μL Invitrogen) in 5 μL of 10X DNase I reaction buffer and 44 μL laboratory grade H2O. Then DNase I was inactivated with 5 μL of 25 mM EDTA (pH 8.0) and 10 min incubation at 65°C. Genomic DNA was extracted and *rpoB* mutant abundance was assessed by RM-seq as described above. Differential abundance analysis of the mutant before and after daptomycin exposure was performed with the R DESeq2 (1.10.1) package (Love et al. 2014, using the count of mutation calculated from table output of the RM-seq data processing pipeline. DESeq2 analysis was performed with all mutations count superior to 1 using default parameters and Cooks cut-off set to false. The Wald statistical test performed by DESeq2 to estimate the significance of the changes in mutation abundance after exposure to daptomycin was used to screen *rpoB* mutations that were associated with increased or decreased tolerance to daptomycin. The detailed explanation of this test is described in (Love et al. 2014). Wald test P values were adjusted for multiple testing using the procedure of Benjamini and Hochberg (Benjamini and Hochberg 1995).

### Mouse infection model

Wild-type 6-week-old female BALB/c mice were injected via the tail vein with approximately 2 × 10^6^ colony-forming units (CFU) in a volume of 100 μL PBS. The mice were monitored every 8 hours until completion of the experiment and were euthanized after 1 day or 7 days post infection. Bacteria from the liver, kidney and spleen were recovered by mechanical homogenization in 1 mL of phosphate buffered saline (PBS), serially diluted and plated on BHI plates. Colonies forming after overnight incubation at 37°C were pooled and assessed by RM-seq. All experiments were performed in accordance with protocols approved by the animal ethics and welfare committee of the University of Melbourne (approval number 1212591).

## DATA ACCESS

Sequencing reads are available from NCBI/ENA/DDBJ under BioProject numbers PRJNA360176 and PRJNA399605. The RM-seq bioinformatics pipeline is available from Github (https://github.com/rguerillot/RM-seq).

## ACKNOWLEDGMENTS

This work was supported by the National Health and Medical Research Council (NHMRC), Australia project grant (GNT1066791) and Research Fellowship to TPS (GNT1008549) and Practitioner Fellowship to BPH (GNT1105905). Doherty Applied Microbial Genomics is funded by the Department of Microbiology and Immunology at The University of Melbourne.

## AUTHOR CONTRIBUTIONS

RG, TPS and BPH designed and planned the project. RG, LL, SB, BH, WG, AD and TT designed and performed the laboratory experiments. RG, TT, IM, SP, SG, AG, MBS, AYP, TPS and BPH provided intellectual input and analysed the data. The manuscript was drafted by RG, TPS and BPH. All authors reviewed and contributed to the final manuscript.

## DISCLOSURE DECLARATION

None to declare

## SUPPLEMENTAL MATERIAL

**Supplemental_Fig_S1: 3D model of S. aureus RpoB in complex with rifampicin.** Rifampicin molecule is coloured in red. The RpoB residue surface a distance of less than 10Â are coloured in white, residues associated with rifampicin resistance by RM-seq are coloured in orange. S. aureus NRS384 WT RpoB protein structure was modelled on the Swiss-model server (https://swissmodel.expasy.org) using Escherichia coli RNA polymerase and rifampicin complex structure (5UAC). The structure model was visualised using PyMOL software.

**Supplemental_Fig_S2:** Prediction of the number of consensus reads at different sequencing depths.

**Supplemental_Table_S1:** Whole genome sequencing and SNP/indel calling of reconstructed rpoB mutants.

**Supplemental_Table_S2:** Rifampicin and daptomycin MICs of reconstructed mutants. **Supplemental_Table_S3:** Primers sequences used for mutations reconstruction and RM-seq.

